# Leukocyte cytokine responses in adult patients with mitochondrial DNA defects

**DOI:** 10.1101/2021.12.13.472449

**Authors:** Kalpita R. Karan, Caroline Trumpff, Marissa Cross, Kristin M. Englestad, Anna L. Marsland, Peter McGuire, Michio Hirano, Martin Picard

## Abstract

Patients with oxidative phosphorylation (OxPhos) defects causing mitochondrial diseases appear particularly vulnerable to infections. Although OxPhos defects modulate cytokine production *in vitro* and in animal models, little is known about how circulating leukocytes of patients with inherited mitochondrial DNA (mtDNA) defects respond to acute immune challenges. In a small cohort of healthy controls (n=21) and patients (n=12) with either the m.3243A>G mutation or single, large-scale mtDNA deletions, we examined: i) cytokine responses (IL-6, TNF-α, IL-1β) in response to acute lipopolysaccharide (LPS) exposure, and ii) sensitivity to the immunosuppressive effects of glucocorticoid signaling (dexamethasone) on cytokine production. In dose-response experiments to determine the half-maximal effective LPS concentration (EC_50_), relative to controls, leukocytes from patients with mtDNA deletions showed 74 - 79% lower responses for IL-6 and IL-1β (p_IL-6_=0.031, p_IL-1β_=0.009). Moreover, IL-6 response to LPS in presence of GC was also blunted in cells from patients with mtDNA deletions (p_IL-6_=0.006), but not in leukocytes from patients with the m.3243A>G mutation. Overall, these *ex vivo* data provide preliminary evidence that some systemic OxPhos defects may compromise immune cytokine responses and glucocorticoid sensitivity. Further work in larger cohorts is needed to define the nature of immune dysregulation in patients with mitochondrial disease, and their potential implications for disease phenotypes.

## Introduction

Mitochondria regulate innate and adaptive immune responses by orchestrating metabolic signals required for immune cell activation, differentiation, and survival [1]. During infections, host immunometabolic responses contribute to normal leukocyte activation [2, 3], such as monocyte polarization into pro- and anti-inflammatory states linked to distinct cytokine secretory profiles [4]. T lymphocyte activation also depends on major bioenergetic recalibrations [5]. Furthermore, we and others have shown that during a targeted pro-inflammatory challenge with the bacterial cell wall molecule lipopolysaccharide (LPS), acutely perturbing mitochondrial oxidative phosphorylation (OxPhos) with pharmacological inhibitors alters cytokine production [6, 7]. This demonstrated in healthy individuals that OxPhos function modulates leukocyte cytokine responses.

In patients with mtDNA disorders, infections are frequent causes of death [8, 9], but what makes adult patients more susceptible or vulnerable to infection remains unclear. One potential factor that may contribute to increased vulnerability to infections is an impaired ability of circulating leukocytes to effectively mount the required innate and adaptive immune responses. During pathogen exposure, the robust activation of immune cells requires effective cell-cell signaling via cytokines [10]. Cytokine responses represent a critical aspect of normal immune function. A systematic review of the literature on mitochondrial diseases confirmed that little is known about cytokine production in affected patients, and that no study has thus far characterized the leukocyte cytokine responses to targeted immune challenges in patients with mtDNA defects.

In vitro, LPS triggers multiple inflammatory cascades that lead to the production of multiple cytokines [11, 12] (reviewed in [13]). This includes the pro-inflammatory cytokines interleukin-6 (IL-6), tumor necrosis factor-alpha (TNF-α), and Interleukin-beta (IL-1β), which are also physiologically elevated by acute physical [14, 15] and psychological stress [16, 17], indicating their broad physiological significance and relative lack of specificity. Moreover, we note that cytokines responses are physiologically regulated by glucocorticoid signaling, which, at nanomolar concentration, potently suppresses pro-inflammatory cytokines, particularly IL-6 [18–20]. Experimentally, applying the glucocorticoid mimetic dexamethasone (Dex) in parallel with LPS to human blood leukocytes therefore allows quantitative assessment of glucocorticoid sensitivity, which reflects an important aspect of immune regulation.

Here, we report LPS-induced pro-inflammatory cytokine responses in blood leukocytes from patients with a pathogenic mtDNA point mutation (m.3243A>G, hereafter “mutation”), or with a single, large-scale mtDNA deletion (hereafter “deletion”). We also measure leukocyte glucocorticoid sensitivity, providing converging preliminary evidence for potential immune alterations in mitochondrial disorders.

## Methods

### Participant recruitment

Informed consent was obtained in compliance with guidelines of the Institutional Review Board of the New York State Psychiatric Institute IRB#7424. All participants provided informed consent for the study procedures and publication of data. Participants between the ages of 18-55 years were recruited between June 2018 and March 2020 as part of the larger Mitochondrial Stress, Brain Imaging, and Epigenetics (MiSBIE) study cohort. The first 21 healthy controls and 12 patients with mitochondrial diseases (Table 1) were included in this sub-study.

**Table1.** Participant characteristics

| Variable <sup>a</sup> | Controls | m.3243A>G carriers | Single, large-scale deletion | *Abbreviations:*<br>BMI, body mass index ;<br>CNS, Columbia Neurological Scale ;<br>NMDAS, Newcastle Mitochondrial Disease |
| --- | --- | --- | --- | --- |
| N | 21 | 7 | 5 |  |
| Age, years | 32.9 (9.5) | 32.1 (9.5) | 33.1(9.9) |  |
| Sex, female | 15 (72) | 3 (42) | 4 (80) |  |
| BMI | 25.8 (6.9) | 24.2 (5.0) | 26.1 (4.1) |  |
| Race/ethnicity |  |  |  |  |
| White | 15 (71.4) | 7 (100) | 5 (100) |  |
| Black | 4 (19) | - | - |  |
| Asian | 1 (4.7) | - | - |  |
| Hispanic or Latino | 1 (4.7) | - | - |  |
| Multiple | - | - | - |  |
| MELAS diagnosis | - | 1 | - |  |
| CPEO diagnosis | - | 1 | 2 |  |
| CPEO “Plus” diagnosis |  |  | 2 |  |
| Multi-systemic syndrome |  | 1 | 1 |  |
| NMDAS score | 1.71 (2.7) | 4.14 (4.0) | 24 (11) |  |
| CNS score | 72 (2.1) | 70.2 (1.3) | 62.25 (7.6) |  |

Assessment Scale.
<sup>a</sup> Data presented as means (standard deviation) or n (%)

Participants were recruited from our local clinic at the Columbia University Irving Medical Center and nationally in the USA and Canada. Patients with mitochondrial diseases were eligible for inclusion if they have a molecularly defined genetic diagnosis for either: i) the m.3243A>G point mutation, with or without mitochondrial encephalopathy, lactic acidosis, and stroke-like episodes (MELAS); or ii) a single, large-scale mtDNA deletion associated chronic progressive external ophthalmoplegia (CPEO) or Kearns-Sayre syndrome (KSS). Exclusion criteria were severe cognitive deficit or inability to provide informed consent, neoplastic disease, symptoms of flu or other seasonal infection four weeks preceding hospital visit, Raynaud’s syndrome, involvement in any therapeutic or exercise trial listed on ClinicalTrials.gov, steroid therapy (e.g., oral dexamethasone, prednisone, or similar), other immunosuppressive treatment, metal inside or outside the body or claustrophobia that would preclude magnetic resonance imaging (MRI). All participants completed a brief questionnaire to collect information on their sex, age, ethnicity, health condition, and medication use. Symptoms severity was measured using the Newcastle Mitochondrial Disease Adult Scale (NMDAS) [21].

### Differential blood counts

Complete blood counts (CBC) were performed on all participants and included proportions of white blood cells (WBC), red blood cells, platelets, and differential WBC counts using an automated hematologic analyzer (XN-9000 Sysmex systems). This yielded absolute cell counts and proportions (%) of total WBC that are neutrophils, lymphocytes, monocytes, eosinophils, and basophils.

### Whole blood LPS-stimulation and glucocorticoid suppression

Whole blood was collected in heparin vacutainer tubes (#BD 367878) and diluted with 1x RPMI without Phenol red (Thermofisher #11835055). For dose-dependent lipopolysaccharide (LPS) stimulation, blood was incubated with bacterial endotoxin LPS from *Escherichia coli* (Sigma-Aldrich, #L2880) at increasing concentrations of 3.2 pg/mL–50 ng/mL per well in a 96-well tissue culture plate (Eppendorf, #30730127). LPS is a potent Toll-like receptor (TLR) 4 agonist derived from outer membrane of Gram-negative bacteria [22, 23]. Relative to isolated cell preparations, the whole blood preparation imposes less stress on leukocytes, preserves physiologically relevant interactions between circulating leukocytes, and preserves the influence of potential circulating humoral factors on immune responses [24].

In glucocorticoid suppression experiments, the cortisol-mimetic dexamethasone (Dex, Sigma-Aldrich, #D4902) was co-incubated at a final concentration of 100 nM [25] with LPS (3.2 pg/mL-50 ng/mL) in whole blood. Each plate consisted of an untreated blood-only control well for baseline measures. The plate was incubated at 37°C with 5% CO_2_ for 6 hrs. Plasma was collected from each well of the plate by centrifuging the plate, first at 1,000g for 5 mins and then at 2,000g for 10 mins at 4°C to isolate cell-free plasma. Plasma was stored at −80°C for cytokine measures.

Three plasma cytokines: interleukin-6 (IL-6), tumor Necrosis Factor-alpha (TNF-α) and interleukin-1beta (IL-1β), were measured using the Abcam Human Catchpoint Simple Step ELISA kits (Abcam, #ab229434, ab229399, and ab229384), a fluorescence-based detection method, following manufacturer’s instructions. Briefly, standards were prepared by serial dilutions to generate 12-14-point standard curves. Plasma samples were diluted to 1:4 ratio using a diluent reagent for all cytokine measures. Fifty microliters of standards and samples were added to appropriate wells in the 96-well strip plate followed by the addition of 50 μL of capture- and detection Ab cocktail to all the wells which was then incubated on a shaker at RT for 1 hr. Post incubation, the wells were aspirated and washed 3x times with 1X wash buffer. After the final wash, a 100 μL of prepared CatchPoint HRP Development Solution was added to the wells and incubated for 10 mins at RT. The plates were read for fluorescence per well at an Ex/Cutoff/Em 530/570/590 nm in in a micro-plate reader (SpectraMax M2, Molecular Devices). Two plasma samples with known cytokine levels were used as internal standards per plate to account for inter-assay variations. A background correction was applied to all RFU values in a run based on ‘no-sample’ blank values. The cytokine concentrations were interpolated from respective standard curves and the final concentration was obtained by correcting for dilution factor.

### LPS-induced cytokine sensitivity

To determine cytokine sensitivity of each participant, a 4-parameter logistic regression was fitted to the absolute circulating levels of IL-6, TNF-α, and IL-1β increasing LPS concentrations (0.0032ng/mL-50ng/mL). Half maximal effective concentrations of LPS (EC_50_) for each cytokine response was derived for each participant using best-fit regression values on the dose-response curve. The maximal cytokine response was obtained at 50ng/mL of LPS (EC_max_) for all participants. Mean EC_50_ and mean max cytokine responses were obtained for both control and disease (mutation and deletion) groups for downstream statistical analyses.

### Statistical analyses

One-way ANOVAs with Dunnett’s multiple comparisons were used to test group differences in LPS- and LPS+Dex -treated cytokine responses. Paired t-tests were used to examine intra-individual differences from pre- to post-Dex treatment. The effect size estimate Hedge’s *g* [26], derived from the variance within groups relative to the group differences, was calculated to obtain a standardized estimate of the magnitude of the effect independent of sample size. Spearman rank correlations were used to quantify the strength of the association between cytokine responses and leukocyte cell counts. All statistical analyses were performed using GraphPad Prism v8.2. p values *p*<0.05 were considered statistically significant.

## Results

### LPS-stimulated cytokine responses

A total of 21 healthy controls and 12 patients (7 3243A>G mutation, 5 single, large-scale deletion) were recruited for this project (see Table 1 for participant characteristics). The mutation group generally presented with mild disease severity (NMDAS score=4.1) whereas patients with deletions were more severely affected (NMDAS=14.0, **Table 1**). Cytokine responses were quantified as: i) cytokine levels at maximal LPS concentration (EC_max_), and ii) the sensitivity of cytokine responses, calculated as the half-maximal effective concentration of LPS required to elicit 50% of the maximal response (EC_50_) (**Figure 1a**).

**Figure 1.**
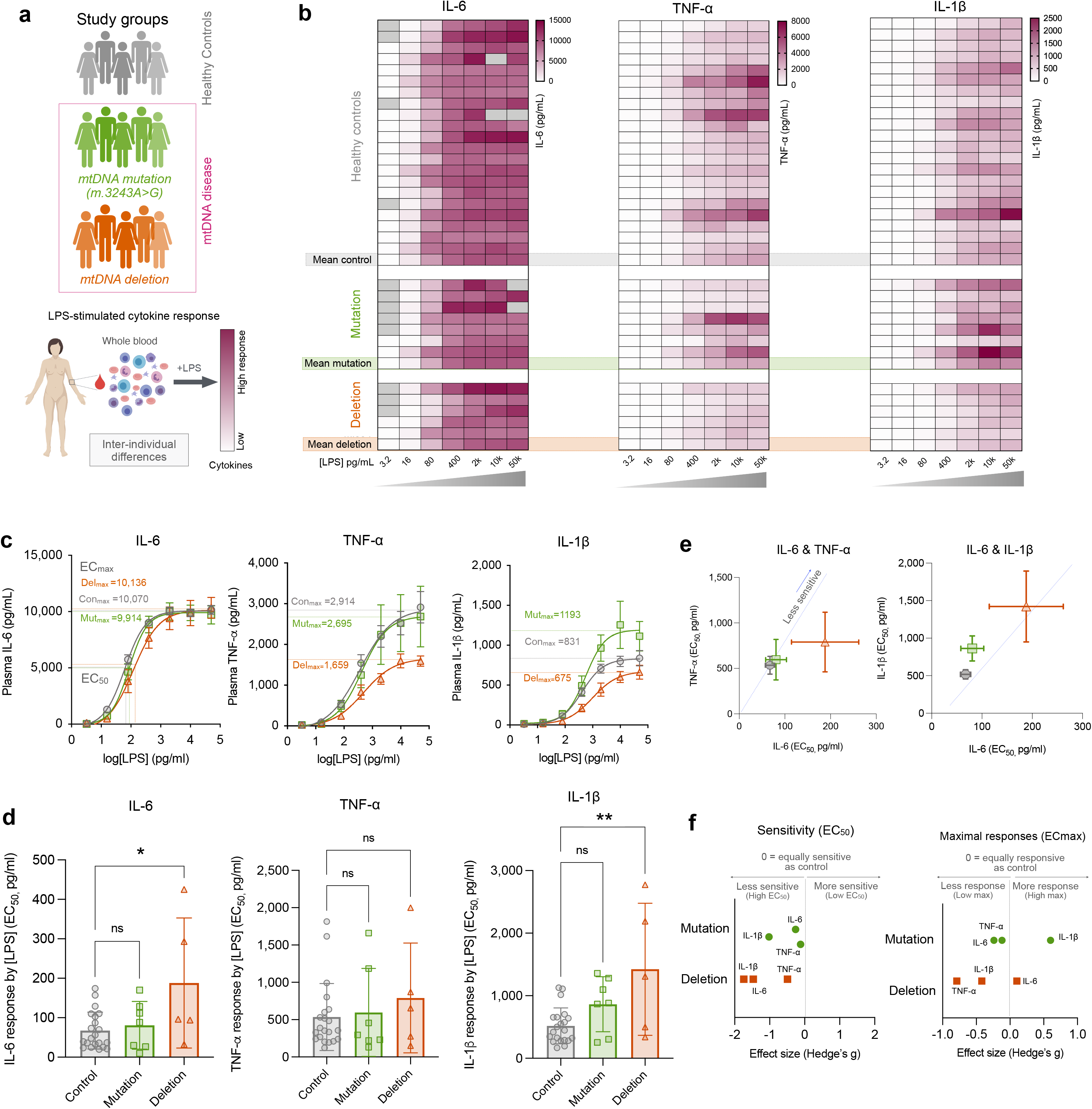
Inter-individual differences in stimulated cytokine responses in patients with mitochondrial disease. a) Schematic of study groups that participated in whole blood LPS exposure. b) Pro-inflammatory cytokine (IL-6, TNF-α and IL-1β) response in whole blood from healthy controls, patients with 3243A>G mutation and single deletions. c) Determination of max (ECmax) and half max conc. of LPS (EC_50_) based on dose-dependent cytokine response to LPS assessed in controls (grey) and patients with mitochondrial disease (green for mutation, and orange for deletion). d) Inter-individual differences in cytokine sensitivity (EC_50_) among patients with mito-disease and healthy controls. e) Cytokine sensitivity bi-plots based on half-max cytokine response (EC_50_). The dotted line passes by the origin and average of the control group. F) Standardized effect sizes (Hedge’s *g*) at 95% confidence interval of cytokine response (EC_50_, *left*; max response, *right*) in disease groups relative to healthy controls. Non-linear regression analysis was used in dose-response curves presented in c, One-way ANOVA with Dunnett’s multiple comparison post-hoc analysis was used in d. *p<0.05 and ** p<0.01; ns, not significant. n=21-22 Controls, Mutation n=7, Deletion n=5.

As expected, IL-6, TNF-α and IL-1β cytokine levels increased substantially (range 30-4,190-fold, ps<0.0001) in response to LPS across all groups (**Figure 1b, c**). There was no significant difference between control and disease groups in the mean IL-6 levels at EC_max_ (one-way ANOVA, p=0.94). However, relative to healthy controls, deletion patients produced 45% less TNF-α (p=0.35) and 21% less IL-1β (p=0.78) at EC_max_ **(Supplemental Figure S1**). The cytokine responses were not associated with the proportions of lymphocytes, monocytes, or neutrophils (**Supplemental Figure S2**).

In relation to LPS sensitivity, consistent with the trend towards reduced peak responses, relative to controls, deletion patients exhibited 1.5- and 2.7-fold higher EC_50_ for TNF-α and IL-1β, respectively (**Figure 1d**). This translates into 48% and 74% lower leukocyte sensitivities for TNF-α and IL-1β, respectively, meaning that deletion patients required higher LPS doses to elicit comparable TNF-α and IL-1β responses to controls. The diverging cytokine response phenotypes among groups are illustrated in bi-variate plots in **Figure 1e**.

Because the small sample sizes increased the probability that a true difference may be missed (i.e., false negatives), we also computed standardized effect sizes (Hedges’ *g*) [26] of the maximal responses and EC_50_-based sensitivity (**Figure 1f)**. These results indicate that blood leukocytes from deletion patients when compared with the control group exhibit blunted sensitivity of moderate (g>0.5) to large (g>0.8) effect sizes, whereas potential alterations in mutations patients are generally in the same direction, but small-to-moderate in magnitude.

### Glucocorticoid sensitivity

Resistance to glucocorticoid-mediated cytokine suppression indicates impaired immune regulation (reviewed in [20, 27]). To examine glucocorticoid sensitivity in mitochondrial diseases, we repeated the LPS dose-response curves in the presence of the glucocorticoid receptor agonist dexamethasone (Dex) (100nM) (**Figure 2a**). All study groups showed the expected anti-inflammatory response to GC administration, illustrated by decreased IL-6, TNF-α and IL-1β responses at EC_max_ (**Figure 2b**). **Supplementary Figure S3** shows the paired pre- and post-Dex individual-level cytokine levels.

**Figure 2.**
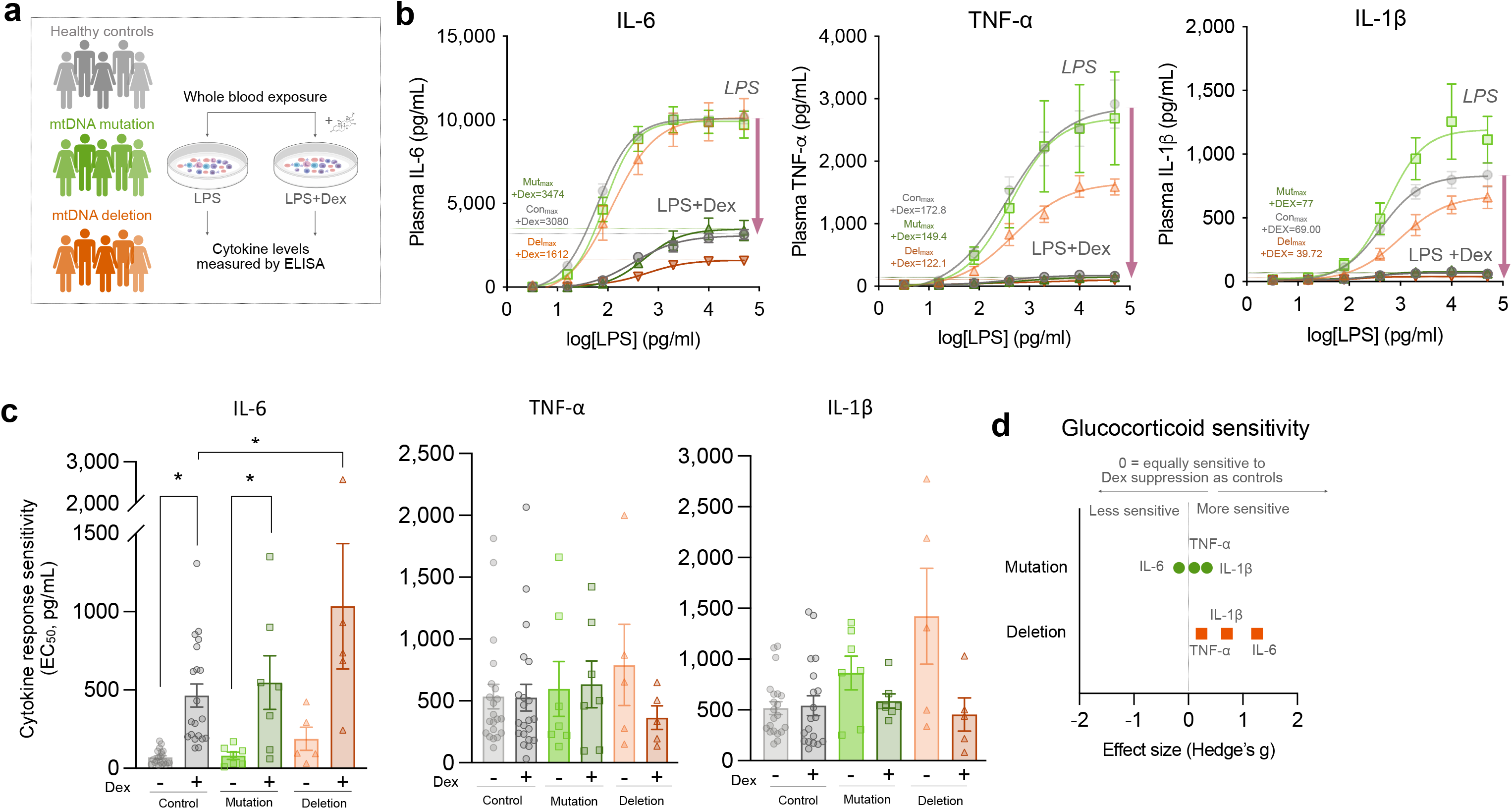
Glucocorticoid sensitivity to LPS-stimulated cytokine responses in mitochondrial diseases. a) Overview of the dexamethasone (Dex) sensitivity assay in whole blood. b) LPS dose-response curves for IL-6, TNF-α and IL-1β without Dex (*top traces*, same as in Figure 1) and with Dex (*bottom traces*). The dotted lines mark the maximal cytokine responses (EC_max_) in healthy participants (*grey*) and patients with the m.3243A>G mutation (*green*) and single, large scale mtDNA deletions (*orange*). c) Effect of Dex on EC_50_ LPS sensitivity; note that higher EC50 values represent lower sensitivity (greater dose required to elicit half of the maximal cytokine response). d) Summary of effect sizes (Hedge’*g)* for deletion and mutation groups relative to control. g>0.5 is a medium effect size, g>0.8 is a large effect size. One-way ANOVA with Dunnett’s multiple comparison test was used in c to test for group differences. Pre- to post-Dex differences were tested with paired t-tests. *p<0.05. n=21-22 Controls, Mutation n=7, Deletion n=5.

Compared to controls, the leukocytes of deletion patients were more sensitive to the glucocorticoid suppression of IL-6 levels. Compared to the 3.8-fold increase in EC_50_ with Dex in controls, Dex increased EC_50_ by 6.6-fold in the deletion group (p=0.006, **Figure 2c**). Dex did not significantly alter EC_50_ for TNF-α or IL-1β, indicating that the sensitization to glucocorticoid signaling was cytokine specific. The effects sizes for Dex-induced cytokine suppression (**Figure 2d**) were small (g>0.1) to moderate (g>0.5) in deletion patients and mostly small (g>0.2) in mutation group. Further, relative to controls, the Dex-induced suppression of EC_max_ IL-6 levels was 12.7% more potent in deletion patients (p=0.026) (**Supplemental Figure S3c**), again indicating that leukocyte IL-6 production is more easily suppressed (i.e., less robust) in the blood of deletion patients. No significant effects were observed in the 3243A>G mutation group.

## Discussion

In this study, we have quantified LPS-induced leukocyte cytokine responses and glucocorticoid sensitivity in the blood of patients with two different primary mitochondrial DNA defects. Patients with single, large-scale mtDNA deletions showed significantly reduced pro-inflammatory IL-6 and IL-1β sensitivity to LPS, along with an exaggerated glucocorticoid-mediated IL-6 suppression, indicating a less robust cytokine production capacity. These preliminary results in a small patient cohort converge to suggest that subgroups of patients with mtDNA defects may exhibit deficient cytokine production capacity and regulation. If confirmed in larger studies, the blunted leukocyte cytokine responses to an acute immune agonist could contribute to poor immune responses, and thus to the vulnerability of patients with mitochondrial diseases to infectious disorders [8, 28].

The mechanism for the blunted cytokine response remains unclear. Using blood from healthy individuals, we previously showed that acutely inhibiting individual OxPhos complexes with specific inhibitors reduced both maximal cytokine levels and sensitivity (EC_50_) to LPS [7]. These pharmacological findings align with our current results in mitochondrial disease, where OxPhos dysfunction is instead of genetic origin. Our previous acute pharmacological design where OxPhos inhibition was initiated at the same time as the LPS challenge implicates direct intracellular processes, such as energy deficiency, redox alterations, or other intracellular signal in this response. However, in patient leukocytes where the mtDNA defect exists since development, or where mtDNA defects trigger systemic, cell non-autonomous effects that influence leukocytes, we cannot rule out other potential factors such as reduced receptor expression (i.e., toll-like receptor 4, for LPS), variable kinetics of cytokine production, or other humoral factors that may exert secondary effects on immune cells [29]. Additional work is necessary to understand the basis for these effects.

An essential aspect of immune function is the regulation of cytokines by secondary neuroendocrine factors, such as glucocorticoids [30]. Natural differences in GC-sensitivity attributable to glucocorticoid receptor subtypes have been previously documented [31], and clinically may underlie the resistance to anti-inflammatory glucocorticoid treatment – i.e., glucocorticoid resistance [32]. To our knowledge, immune glucocorticoid sensitivity has not previously been investigated in mitochondrial diseases. Our finding that deletion patients are more sensitive to GC-induced IL-6 suppression suggests the opposite to glucocorticoid resistance, possibly a form of glucocorticoid “hypersensitivity” as it relates to IL-6 cytokine down-regulation. An earlier report showed that immunosuppressed mice with elevated corticosterone partially recovered by restoring the energy balance using pyruvate [33]. This mitochondrial regulation of glucocorticoid sensitivity is also in line with previous work where we showed that inhibiting complex I similarly increased leukocyte GC sensitivity by 12.3% (Dex-mediated IL-6 suppression) [7]. Thus, in the absence of healthy OxPhos system, or possibly to secondary systemic signals of OxPhos dysfunction, stimulated leukocyte cytokine production is less robust, and more easily suppressed. If these effects are confirmed, increased glucocorticoid sensitivity in subgroups of patients with mitochondrial diseases could have implications for their clinical care, possibly warranting additional caution for the use of steroid therapy in this population. This result could also have implications for patients’ resilience towards psychosocial stressors that stimulate the hypothalamic-pituitary-adrenal (HPA) axis, resulting in the secretion of the endogenous immunosuppressive glucocorticoid hormone cortisol [34].

Some limitation of this study should be noted. Although of small sample size, a strength of our study is the homogeneity of our disease groups, which either harbor the same mtDNA point mutation (m.3243A>G) or a single, large-scale mtDNA deletion, established by clinical genetic testing. Leukocyte heteroplasmy could not be determined in this study and represents a major factor that future studies need to address in detail. However, we note that large-scale mtDNA deletions are depleted from immune cells lineages, and thus rarely detectable in blood [35–37] possibly due to purifying selection in blood leukocytes and bone marrow [38]. In comparison, the m.3243A>G mutation and other point mutations are typically present in immune cells [39] and may segregate between different immune lineages [40]. Hence, our main observations regarding blunted cytokine response in circulating leukocytes of deletion patients is particularly intriguing. The specificity of this finding for deletion, and not 3243A>G carriers could suggest cell non-autonomous effects mediated by unknown humoral factors arising from somatic tissues that harbor mtDNA deletions [35, 41–43]. We also note that the greater disease severity of the deletion group could contribute to explain the group differences, independently from the specific genetic defects. Although no patient was on steroid hormone, we also cannot rule out the possibility that medications that may influence cytokine responses, which our sample did not allow to control.

In summary, our *ex vivo* results in a small cohort of patients with two different mtDNA defects provide preliminary evidence that immune sensitivity and cytokine responses are blunted in patients with single, large-scale mtDNA deletions. This work provides initial evidence in need of validation in larger studies, and if confirmed, could help to understand why adult patients with mitochondrial diseases are at increased risk to die of infectious conditions.

## Statements & Declarations

### Author contributions

K.R.K. and M.P. designed the study. C.T., M.H., M.P. developed the clinical protocol. K.E. and M.H. provided clinical diagnosis and recruited patients. M.C. coordinated study activities and collected data. K.R.K. performed LPS stimulation and cytokine measurements. K.R.K. analyzed data and prepared the figures. K.R.K. and M.P. drafted the manuscript with P.M. All authors reviewed and edited the final version of this manuscript.

## Acknowledgements

We thank the patients and healthy volunteers who contributed to the study, Johanne Fortune for assistance with phlebotomy, and the rest of the MiSBIE Team. We acknowledge Logan Beharry for his contribution to the systematic review of the literature.

## Data availability

Requests for additional information and data will be fulfilled by the corresponding author.

## Consent to participate and publish

Informed consent was obtained in compliance with guidelines of the Institutional Review Board of the New York State Psychiatric Institute IRB#7424. All participants provided informed consent for the study procedures and publication of data.

---

**Supplementary Figure S1. Cytokine response to increasing LPS exposure in whole blood from patients with mitochondrial disease and healthy controls.** Represented are IL-6, TNF-α and IL-1β levels in response to increasing level of exposure to LPS in patients with mito-disease (Mutation in green, deletion in orange) and healthy controls (grey). n=21-22 Controls, Mutation n=7, Deletion n=5.

**Supplementary Figure S2. Correlation between proportions of leukocytes (%) and whole blood cytokine responses.** a-c) Proportions of leukocyte subtypes were obtained from complete differential counts (CBC) data, n=33 (all participants including controls and mito disease patients). Spearman rho (r) correlation analysis was performed between % leukocyte types and individual cytokine responses (EC_50_ presented here). The grey lines represent simple linear regressions and grey shaded regions show 95% confidence intervals. p<0.05 is significant.

**Supplementary Figure S3. GC-sensitivity of cytokines in patients with mitochondrial disease.** a) Experiment design to evaluate cytokine response pre- to post-Dex among the participating groups: controls, patients with mutation and patients with large deletion. b) Dex-sensitivity of cytokines in participants. Within-group comparisons of pre- to post-Dex cytokine responses at half-max LPS (EC_50)._ c) Average % suppression of cytokine levels by Dex (GC-sensitivity) of patients and controls. Paired t-test was performed in b to test significant Dex effects within groups and One-way ANOVA was performed in c to test significant (mean) group difference. *p<0.05 is significant.

## References

1. Angajala, A., et al., Diverse Roles of Mitochondria in Immune Responses: Novel Insights Into Immuno-Metabolism. Front Immunol, 2018. 9: p. 1605.

2. Shi, L., et al., Biphasic Dynamics of Macrophage Immunometabolism during Mycobacterium tuberculosis Infection. mBio, 2019. 10(2).

3. Gleeson, L.E., et al., Cutting Edge: Mycobacterium tuberculosis Induces Aerobic Glycolysis in Human Alveolar Macrophages That Is Required for Control of Intracellular Bacillary Replication. J Immunol, 2016. 196(6): p. 2444–9.

4. Van den Bossche, J., et al., Mitochondrial Dysfunction Prevents Repolarization of Inflammatory Macrophages. Cell Rep, 2016. 17(3): p. 684–696.

5. Dumitru, C., A.M. Kabat, and K.J. Maloy, Metabolic Adaptations of CD4(+) T Cells in Inflammatory Disease. Front Immunol, 2018. 9: p. 540.

6. Weiss, S.L., et al., Mitochondrial Dysfunction Is Associated with an Immune Paralysis Phenotype in Pediatric Sepsis. Shock, 2019.

7. Karan, K.R., et al., Mitochondrial respiratory capacity modulates LPS-induced inflammatory signatures in human blood. Brain, Behavior, & Immunity - Health, 2020. 5.

8. Walker, M.A., et al., Predisposition to infection and SIRS in mitochondrial disorders: 8 years’ experience in an academic center. J Allergy Clin Immunol Pract, 2014. 2(4): p. 465–468, 468 e1.

9. Barends, M., et al., Causes of Death in Adults with Mitochondrial Disease, in JIMD Reports, Volume 26, E. Morava, et al., Editors. 2016, Springer Berlin Heidelberg: Berlin, Heidelberg. p. 103–113.

10. Albrecht, L.J., et al., Lack of Proinflammatory Cytokine Interleukin-6 or Tumor Necrosis Factor Receptor-1 Results in a Failure of the Innate Immune Response after Bacterial Meningitis. Mediators Inflamm, 2016. 2016: p. 7678542.

11. Manderson, A.P., et al., Subcompartments of the macrophage recycling endosome direct the differential secretion of IL-6 and TNFalpha. J Cell Biol, 2007. 178(1): p. 57–69.

12. Netea, M.G., et al., Differential requirement for the activation of the inflammasome for processing and release of IL-1beta in monocytes and macrophages. Blood, 2009. 113(10): p. 2324–35.

13. Kany, S., J.T. Vollrath, and B. Relja, Cytokines in Inflammatory Disease. Int J Mol Sci, 2019. 20(23).

14. Domin, R., et al., Effect of Various Exercise Regimens on Selected Exercise-Induced Cytokines in Healthy People. Int J Environ Res Public Health, 2021. 18(3).

15. Hamer, M. and A. Steptoe, Association between physical fitness, parasympathetic control, and proinflammatory responses to mental stress. Psychosom Med, 2007. 69(7): p. 660–6.

16. Marsland, A.L., et al., The effects of acute psychological stress on circulating and stimulated inflammatory markers: A systematic review and meta-analysis. Brain Behav Immun, 2017. 64: p. 208–219.

17. Steptoe, A., M. Hamer, and Y. Chida, The effects of acute psychological stress on circulating inflammatory factors in humans: a review and meta-analysis. Brain Behav Immun, 2007. 21(7): p. 901–12.

18. Bhattacharyya, S., et al., Macrophage glucocorticoid receptors regulate Toll-like receptor 4-mediated inflammatory responses by selective inhibition of p38 MAP kinase. Blood, 2007. 109(10): p. 4313–9.

19. Horton, D.L. and D.G. Remick, Delayed addition of glucocorticoids selectively suppresses cytokine production in stimulated human whole blood. Clin Vaccine Immunol, 2010. 17(6): p. 979–85.

20. Quax, R.A., et al., Glucocorticoid sensitivity in health and disease. Nat Rev Endocrinol, 2013. 9(11): p. 670–86.

21. Schaefer, A., et al., Mitochondrial disease in adults: a scale to monitor progression and treatment. Neurology, 2006. 66(12): p. 1932–1934.

22. Chow, J.C., et al., Toll-like receptor-4 mediates lipopolysaccharide-induced signal transduction. J Biol Chem, 1999. 274(16): p. 10689–92.

23. Pugin J, S.-M.C., Leturcq D, Moriarty A, Ulevitch RJ, Tobias PS., Lipopolysaccharide activation of human endothelial and epithelial cells is mediated by lipopolysaccharide-binding protein and soluble CD14. Proc Natl Acad Sci USA, 1993. 90: p. 2744–2748.

24. Strahler, J., N. Rohleder, and J.M. Wolf, Acute psychosocial stress induces differential short-term changes in catecholamine sensitivity of stimulated inflammatory cytokine production. Brain Behav Immun, 2015. 43: p. 139–48.

25. Alm, J.J., et al., Transient 100 nM dexamethasone treatment reduces inter- and intraindividual variations in osteoblastic differentiation of bone marrow-derived human mesenchymal stem cells. Tissue Eng Part C Methods, 2012. 18(9): p. 658–66.

26. Hedges LV, O., I, Statistical Methods in Meta-Analysis.. 1st ed. 1985: Academic Press Inc.

27. Peter J Barnes, I.M.A., Glucocorticoid resistance in inflammatory diseases. The Lancet, 2009. 373.

28. Pickett, S.J., et al., Phenotypic heterogeneity in m.3243A>G mitochondrial disease: The role of nuclear factors. Ann Clin Transl Neurol, 2018. 5(3): p. 333–345.

29. Vattemi, G., et al., Overexpression of TNF-alpha in mitochondrial diseases caused by mutations in mtDNA: evidence for signaling through its receptors on mitochondria. Free Radic Biol Med, 2013. 63: p. 108–14.

30. Pace, T.W., F. Hu, and A.H. Miller, Cytokine-effects on glucocorticoid receptor function: relevance to glucocorticoid resistance and the pathophysiology and treatment of major depression. Brain Behav Immun, 2007. 21(1): p. 9–19.

31. King, E.M., et al., Glucocorticoid repression of inflammatory gene expression shows differential responsiveness by transactivation- and transrepression-dependent mechanisms. PLoS One, 2013. 8(1): p. e53936.

32. Yang, N., D.W. Ray, and L.C. Matthews, Current concepts in glucocorticoid resistance. Steroids, 2012. 77(11): p. 1041–9.

33. Neigh, G.N., et al., Pyruvate prevents restraint-induced immunosuppression via alterations in glucocorticoid responses. Endocrinology, 2004. 145(9): p. 4309–19.

34. Russell, G. and S. Lightman, The human stress response. Nat Rev Endocrinol, 2019. 15(9): p. 525–534.

35. Jeppesen, T.D., M. Duno, and J. Vissing, Mutation Load of Single, Large-Scale Deletions of mtDNA in Mitotic and Postmitotic Tissues. Front Genet, 2020. 11: p. 547638.

36. Lee, H.F., et al., The neurological evolution of Pearson syndrome: case report and literature review. Eur J Paediatr Neurol, 2007. 11(4): p. 208–14.

37. Trifunov, S., et al., Clonal expansion of mtDNA deletions: different disease models assessed by digital droplet PCR in single muscle cells. Sci Rep, 2018. 8(1): p. 11682.

38. Palozzi, J.M., S.P. Jeedigunta, and T.R. Hurd, Mitochondrial DNA Purifying Selection in Mammals and Invertebrates. J Mol Biol, 2018. 430(24): p. 4834–4848.

39. Grady, J.P., et al., Disease progression in patients with single, large-scale mitochondrial DNA deletions. Brain, 2014. 137(Pt 2): p. 323–34.

40. Walker, M.A., et al., Purifying Selection against Pathogenic Mitochondrial DNA in Human T Cells. N Engl J Med, 2020. 383(16): p. 1556–1563.

41. Forsström S, J.C., Carroll CJ, Kuronen M, Pirinen E, Pradhan S, Marmyleva A, Auranen M, Kleine IM, Khan NA, Roivainen A, Marjamäki P, Liljenbäck H, Wang L, Battersby BJ, Richter U, Velagapudi V, Nikkanen J, Euro L, Suomalainen A., Fibroblast Growth Factor 21 Drives Dynamics of Local and Systemic Stress Responses in Mitochondrial Myopathy with mtDNA Deletions.. Cell Metab, 2019. 30: p. 1040–1054.

42. Lehtonen JM, F.S., Bottani E, Viscomi C, Baris OR, Isoniemi H, Höckerstedt K, Österlund P, Hurme M, Jylhävä J, Leppä S, Markkula R, Heliö T, Mombelli G, Uusimaa J, Laaksonen R, Laaksovirta H, Auranen M, Zeviani M, Smeitink J, Wiesner RJ, Nakada K, Isohanni P, Suomalainen A., FGF21 is a biomarker for mitochondrial translation and mtDNA maintenance disorders. Neurology, 2016. 29(87): p. 2290–2299.

43. Sharma, R., et al., Circulating markers of NADH-reductive stress correlate with mitochondrial disease severity. J Clin Invest, 2021. 131(2).

