## Supplementary Figures for "Leukocyte cytokine responses in adult patients with mitochondrial DNA defects"

### Figure S1

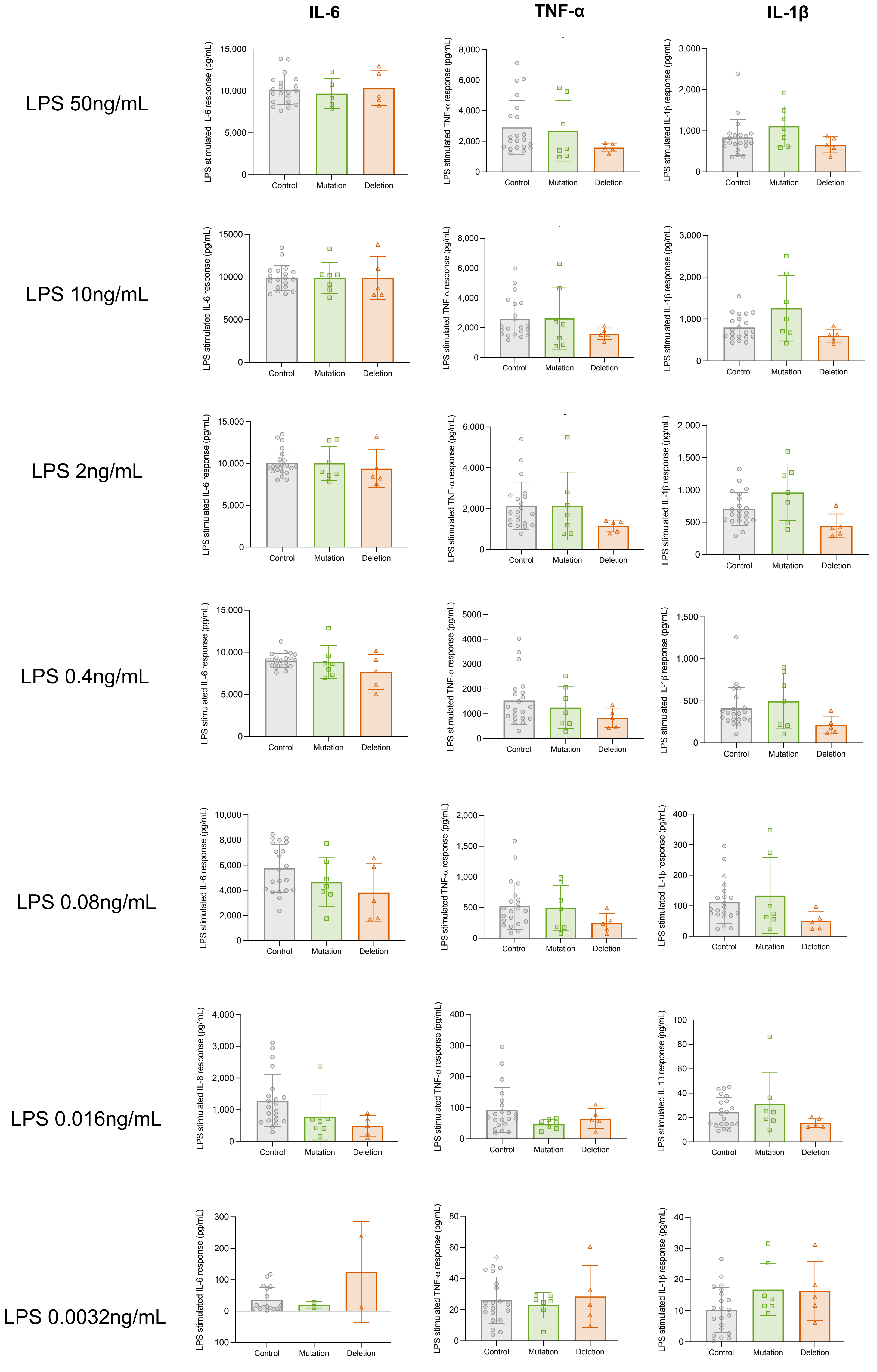


**Figure S1. Cytokine response to increasing LPS exposure in whole blood from patients with mitochondrial disease and healthy controls.** A) Represented are IL-6, TNF-α and IL-1β levels in response to increasing level of exposure to LPS in patients with mito-disease (Mutation in green, deletion in orange) and healthy controls (grey). n=21-22 Controls, Mutation n=7, Deletion n=5.

### Figure S2

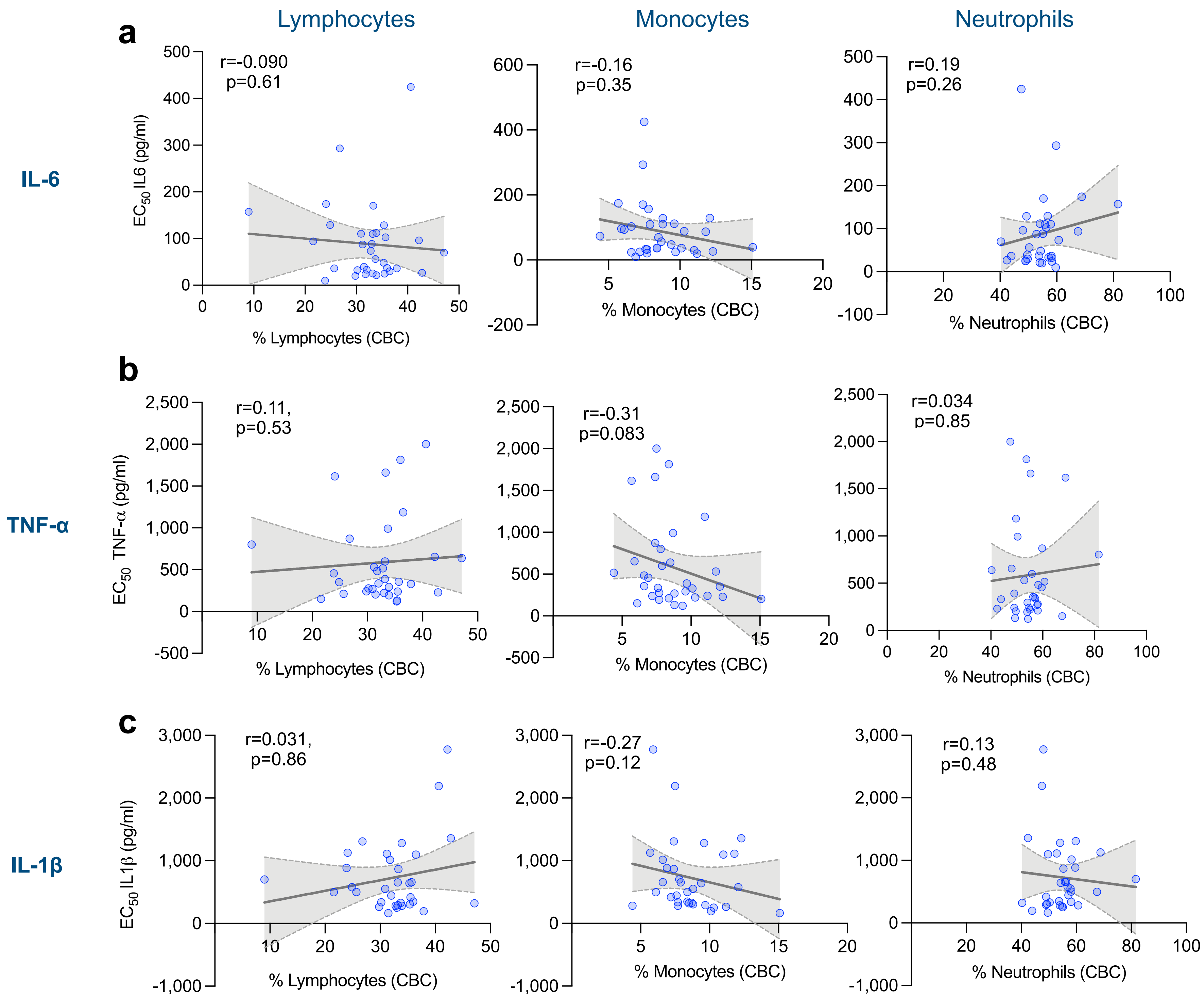


**Figure S2. Correlation between proportions of leukocytes (%) and whole blood cytokine responses.** a-c) Proportions of leukocyte subtypes were obtained from complete differential counts (CBC) data, n=33 (all participants including controls and mito disease patients). Spearman rho (r) correlation analysis was performed between % leukocyte types and individual cytokine responses ($EC_{50}$ presented here). The grey lines represent simple linear regressions and grey shaded regions show 95% confidence intervals. $p<0.05$ is significant.

#### Figure S3

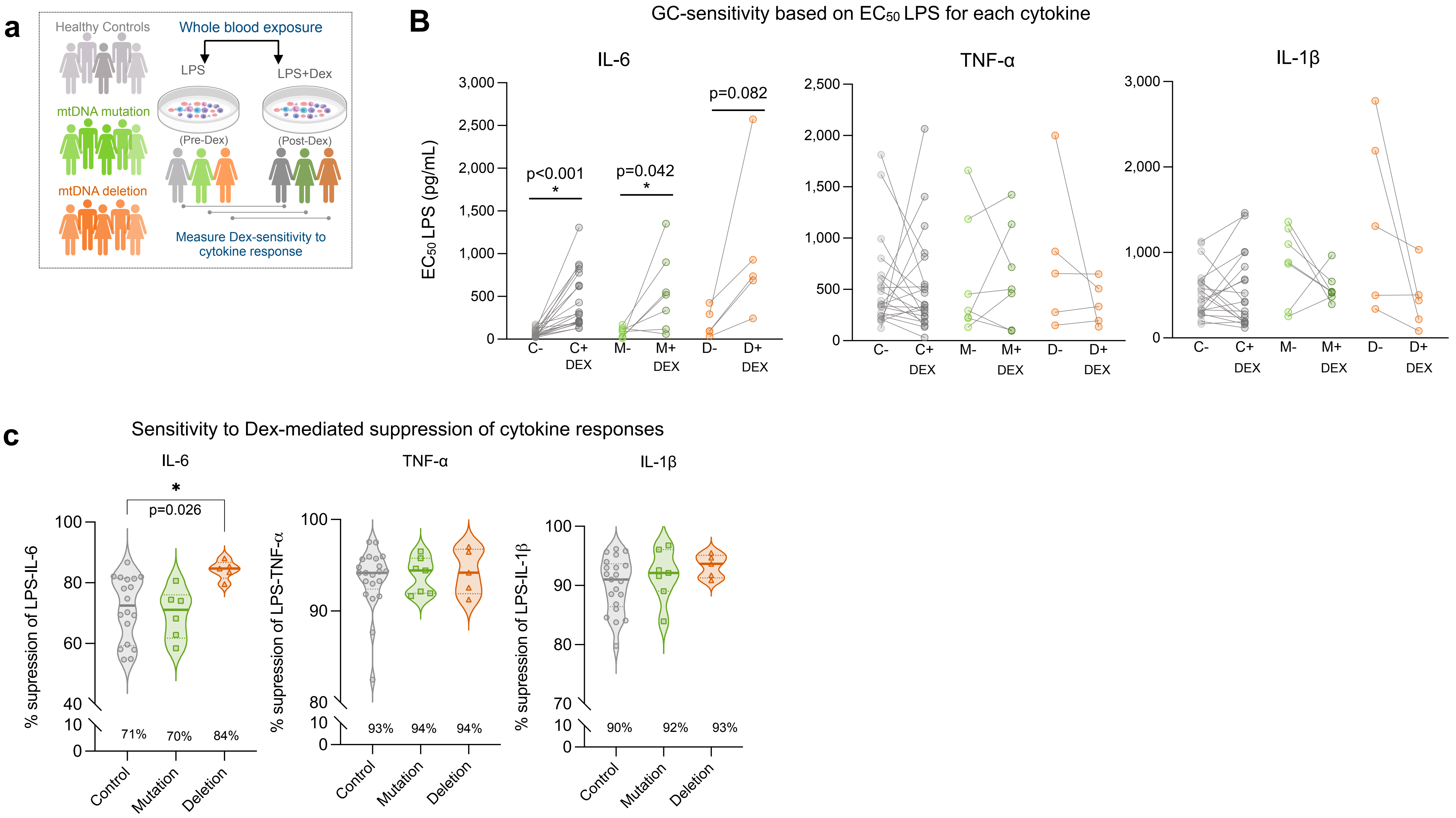


**Figure S3. GC-sensitivity of cytokines in patients with mitochondrial disease.** a) Experiment design to evaluate cytokine response pre- to post-Dex among the participating groups: controls, patients with mutation and patients with large deletion. b) Dex- sensitivity of cytokines in participants. Within-group comparisons of pre- to post- Dex cytokine responses at half-max LPS ($EC_{50}$). c) Average % suppression of cytokine levels by Dex (GC-sensitivity) of patients and controls. Paired t-test was performed in B to test significant Dex effects within groups and One-way ANOVA was performed in 'c' to test significant (mean) group difference. *p<0.05 is significant.
